# Direct antiviral activity of interferon stimulated genes is responsible for resistance to paramyxoviruses in ISG15-deficient cells

**DOI:** 10.1101/2019.12.12.873919

**Authors:** David Holthaus, Andri Vasou, Connor G. G. Bamford, Jelena Andrejeva, Christina Paulus, Richard E. Randall, John McLauchlan, David J. Hughes

## Abstract

Interferons (IFNs), produced during viral infections, induce the expression of hundreds of IFN-stimulated genes (ISGs). Some ISGs have specific antiviral activity while others regulate the cellular response. Besides functioning as an antiviral effector, IFN-stimulated gene 15 (ISG15) is a negative regulator of IFN signalling and inherited ISG15-deficiency leads to autoinflammatory interferonopathies where individuals exhibit elevated ISG expression in the absence of pathogenic infection. We have recapitulated these effects in cultured human A549-ISG15^-/-^ cells and (using A549-UBA7^-/-^ cells) confirmed that posttranslational modification by ISG15 (ISGylation) is not required for regulation of the type-I IFN response. ISG15-deficient cells pre-treated with IFN-α were resistant to paramyxovirus infection. We also showed that IFN-α treatment of ISG15-deficient cells led to significant inhibition of global protein synthesis leading us to ask whether resistance was due to the direct antiviral activity of ISGs or whether cells were non-permissive due to translation defects. We took advantage of the knowledge that IFN-induced protein with tetratricopeptide repeats 1 (IFIT1) is the principal antiviral ISG for parainfluenza virus 5 (PIV5). Knockdown of IFIT1 restored PIV5 infection in IFN-α-pre-treated ISG15-deficient cells, confirming that resistance was due to the direct antiviral activity of the IFN response. However, resistance could be induced if cells were pre-treated with IFN-α for longer times, presumably due to inhibition of protein synthesis. These data show that the cause of virus resistance is two-fold; ISG15-deficiency leads to the ‘early’ over-expression of specific antiviral ISGs, but the later response is dominated by an unanticipated, ISG15- dependent, loss of translational control.

**Key points:** Cell culture model of ISG15-deficiency replicate findings in ISG15^-/-^ patient cells

Cause of resistance in ISG15^-/-^ cells differs depending on duration of IFN treatment

ISG15^-/-^ patients without serious viral disease don’t prove ISGylation is unimportant

## Introduction

The innate immune response against pathogens is underpinned by the evolutionary conserved interferon (IFN) system. All cells express pathogen recognition receptors (PRRs) that sense the products of infection and establish a signalling cascade leading to the production of cytokines, including type I IFN (IFN-α/β) (1, 2). IFN is secreted from cells and binds to cell surface receptors expressed on both infected and non-infected cells, initiating a Janus kinase/signal transducer and activator of transcription (JAK/STAT) signalling cascade, culminating in the expression of hundreds of interferon stimulated genes (ISGs) (3). The biological effects of ISGs are extensive and their principle role is to generate an unfavourable environment for the replication of viruses. Many ISGs have broad antiviral activity, such as double-stranded RNA dependent protein kinase (PKR) that, upon recognition of viral dsRNA, dampens general protein synthesis and prevents the translation of viral mRNAs (4). Other antiviral ISGs, such as IFN-induced protein with tetratricopeptide repeats (IFIT) proteins, inhibit specific viruses, but for many, they are inconsequential (5). Additionally, multiple ISGs are generally required to limit infection because the majority of ISGs result in low to moderate levels of inhibition (6); however, ISGs with specific antiviral properties for a given virus are often not known. Nevertheless, the nature of the innate immune response necessitates the production of the complete spectrum of ISGs, albeit with a high degree of redundancy, as during a natural infection, the identity of the infecting virus is not known. This response is inevitably tightly regulated, as a dysregulated response leads to a suite of autoinflammatory diseases (7).

The ubiquitin-like protein (Ubl) ISG15 is strongly induced by IFN and is critical for regulating how cells respond to infection. As a posttranslational modification (PTM), it can covalently modify proteins in a process known as ISGylation, and in many cases, modification of viral proteins forms part of the antiviral response (8). Covalently bound ISG15 can also be removed from proteins by the ubiquitin specific protease 18 (USP18) (9). Importantly, loss-of-function mutations in ISG15 have been identified in human patients with subsets of autoinflammatory interferonopathies and typically these individuals demonstrate elevated ISG expression in the absence of pathogenic infection (10). Mechanistically, it was shown that ISG15 functions as a negative regulator of type I IFN signalling by stabilising USP18, a known inhibitor of JAK/STAT signalling (11-13). Intriguingly, despite the known functions of ISG15 and USP18 in the ISGylation process, the regulation of type I IFN signalling was entirely independent of ISGylation (10). Interestingly, mouse Isg15 is not required to stabilise Usp18 and appears not to be needed to regulate IFN signalling, suggesting a species-specific, gain-of-function for human ISG15 (14).

Previous work has shown that cells from ISG15-deficient patients expressed higher levels of ISGs compared to normal controls when treated with recombinant IFN-α and these cells were resistant to several viruses (14); however, it was not clear at what stage of infection viruses were blocked nor how. Furthermore, cells were treated with IFN-α followed by washing (to remove IFN) and rested for 36 hours prior to infection. Since ISG15 is involved in regulating the cell cycle (15) and protein synthesis (shown in this report), an over-amplified IFN response (due to lack of ISG15 and reduced levels of USP18) may have led to virus resistance simply because cells were no longer permissive to infection. This has implications for our understanding as to why ISG15-deficient patients are not more susceptible to viral infections; these observations have led to the suggestion that, unlike in mice, human ISG15 is not an antiviral effector (14, 16).

In this study, we recapitulated the phenotype observed in ISG15-deficient patient cells upon treatment with recombinant IFN-α in a cell culture model and dissected the mechanisms that result in virus resistance during an antiviral state. We showed that resistance was due to the direct antiviral activity of the type I IFN response and discuss the implications of ISG15-loss-of-function during the innate immune response. Based on our findings, we conclude that observations from ISG15-deficient patients alone cannot be used to infer that ISG15 does not possess antiviral effector functions, as has been proposed (14, 16).

## Materials and methods

### Cells

Vero cells (African green monkey kidney epithelial cells) and A549 cells (human adenocarcinoma alveolar basal epithelial cells), and derivatives, were maintained in Dulbecco’s modified Eagles’s medium (DMEM; Sigma) supplemented with 10% (v/v) heat-inactivated foetal bovine serum (FBS, Biowest) and incubated in 5% (v/v) CO2 at 37°C in a humidified incubator. A549-shIFIT1 have been described elsewhere (17) and were maintained in blasticidin (10 µg/ml). A549-ISG15^-/-^ cells were generated by CRISPR/Cas9n system that utilises the D10A dual ‘nickase’ mutant of Cas9 (Cas9n) that ostensibly limits off-target effects. Briefly, to disrupt exon 2 of the ISG15 gene, single guide RNA (sgRNA) sequences were cloned using pPX460 and transfected into A549 cells as previously described (18). Transfectants were enriched by treating cells with puromycin (1μg/ml) for 2 d and then diluted to single cells in 96-well plates. Correctly edited cell clones were verified by immunoblot analysis. A549-ISG15^-/-^-shIFIT1 cells were generated as previously described using A549-ISG15^-/-^ (B8) and maintained in media with blasticidin (10 µg/ml) (17). To generate A549-UBA7^-/-^ cells, A549 cells were first made to stably express *Streptococcus pyogenes* Cas9 following blasticidin selection of cells transduced with lentiCas9-Blast (gift from Feng Zhang, Addgene plasmid # 52962 (19)). The sgRNA sequence that targeted exon 3 of UBA7 was chosen computationally (https://www.deskgen.com) and complementary oligonucleotides (sense: caccGCACACGGGTGACATCACTG; antisense: aaacCAGTGATGTCACCCGTGTGC) were hybridised and ligated into the *Bsm* BI site of pLentiGuide-Puro (gift from Feng Zhang, Addgene # 52963 (20)). Cas9-expressing A549s were transduced with UBA7 sgRNA-expressing lentiGuide-Puro and selected with puromycin. Puromycin-resistant cells were single-cell cloned by FACS and successful knockout cells were validated by immunoblot analysis. A549-Npro cells have been described previously (21).

### Virus infections and treatments

Viruses used were human parainfluenza virus 2 (HPIV2) strain Colindale (HPIV2-Co), HPIV3 strain Washington/47885/57 (HPIV3-Wash) (20), PIV5 strain W3 (PIV5-W3) (22) and PIV5 strain CPI- (PIV5- CPI-) (23). Virus stocks were prepared by inoculating Vero cells at a multiplicity of infection (MOI) of 0.001 with continual rocking at 37°C. Supernatants were harvested at 2 d p.i., clarified by centrifugation at 3,000 xg for 15 min, aliquoted and snap frozen. Titres were estimated by standard plaque assay on Vero cells in 6-well plates.

For infection studies, cell monolayers were infected in 6-well plates with virus diluted in medium to achieve a MOI of 10, unless stated otherwise. Virus adsorption was for 1 h, after which the viral inoculum was removed and replaced with media supplemented with 2% (v/v) FBS and incubated in 5% (v/v) CO_2_ at 37°C until harvested. When cells were treated with IFN-α prior to infection (pre-treated) this was done with 1000 IU/ml IFN-α2b (referred to as IFN-α from here on; IntronA, Merck Sharp & Dohme Ltd.) 18 h prior to infection, unless otherwise stated. IFN-α remained on cells for the duration of experiments. Cells were either processed for immunoblot analysis or (if infecting with rPIV5-mCherry, kind gift of Dr He, University of Georgia, USA) imaged using an IncuCyte Zoom imaging system (Sartorius).

For plaque assays 30-40 PFU PIV5-CPI- in 1 ml DMEM, 2% FBS were adsorbed for 1 h onto confluent monolayers of cells in 6-well plates while rocking at 37°C. Following adsorption, 2 ml overlay (DMEM, 2% FBS, Avicel) was added to wells and incubated for 6 d. Cells were fixed with 5% formaldehyde (10 min), washed in PBS and either stained for 10 min with 1 mg/ml toluidine blue O (Sigma) followed by rinses with water or permeabilised for 10 min (PBS, 1% Triton X-100, 3% FBS) washed again and incubated for 1 h with a pool of PIV5-specific antibodies (24) or mouse monoclonal anti-HPIV3 NP (25) diluted in PBS, 3% FBS (1:1000). Following PBS washes, cells were incubated for 1 h with goat anti-mouse IgG antibodies conjugated to alkaline phosphatase (Abcam Cat# ab97020) diluted 1:1000 in PBS, 3%FBS. Cells were washed in PBS and signals were detected using SIGMAFAST BCIP/NBT (Sigma).

### Reverse transcription quantitative PCR

To quantify ISG expression, total cellular RNA was purified from cells that had been treated with 1000 IU/ml IFN-α for 18 h, or left untreated, using TRIzol reagent (ThermoFisher Scientific) and Direct-zol RNA Miniprep Plus kits followed by on-column DNase I treatment for the removal of contaminating DNA (Zymo Research). To measure PIV5-W3 transcription, the indicated cells were treated with 1000 IU/ml IFN-α2b for 8 h and then infected with PIV5 (MOI 10). Following adsorption for 1 h at 37°C, cells were lysed in TRIzol at the indicated times and RNA was purified as above. Complementary DNA (cDNA) was synthesised in 20 µl reaction volumes with 500 ng (ISGs) or 100 ng (PIV5-infected cells) total RNA and oligo(dT) using GoScript reverse transcriptase (Promega) according to the manufacturer’s recommendations. Quantitative PCR reaction mixes (20 μl) included 1x PerfeCTa SYBR green SuperMix (Quanta BioScience), 0.5 μM each primer and 1 μl cDNA reaction mix. Cycling was performed in a Mx3005P real time PCR machine (Stratagene) and included an initial 3 min enzyme activation step at 95°C, followed by 40 cycles of 10 s at 95°C, 10 s at 58°C and 20 s at 72°C. Melting curve analysis was performed to verify amplicon specificity. Quantification of *β-ACTIN* mRNA was used to normalize between samples and the average cycle threshold (CT) was determined from three independent cDNA samples from independent cultures. Relative expression compared to non-treated control cells was calculated using the ΔΔCT method. Primer sequences were: *HERC5* 5’GACGAACTCTTGCACCGTCTC and 5’GCGTCCACAGTCATTTTCCAC, *USP18* 5’ATGCTGCCCAACTGTACCTC and 5’CCTGCAGTCTCTCCACCAAG, *MxA* 5’GCCTGCTGACATTGGGTATAA and 5’CCCTGAAATATGGGTGGTTCTC, *IFIT1* 5’CCTGGAGTACTATGAGCGGGC and 5’TGGGTGCCTAAGGACCTTGTC, PIV5 *NP* 5’AGGGTAGAGATCGATGGCT and 5’GTCTGACCACCATTCCCTT, *β-ACTIN* 5’AGCGAGCATCCCCCAAAGTT and 5’AGGGCACGAAGGCTCATCATT.

### Immunoblotting

Confluent monolayers in 6-well dishes were lysed with 250 µl 2 x Laemmli sample buffer (4% w/v SDS, 20% v/v glycerol, 0.004% w/v bromophenol blue and 0.125 M Tris-HCl, pH 6.8 with 10% v/v β-mercaptoethanol) for 10 min, incubated at 95°C for 10 min, sonicated at 4°C with 3 cycles of 30 s on 30 s off in a Bioruptor Pico (Diagenode) and clarified by centrifugation at 12,000 xg, 4°C for 10 min. SDS-PAGE in Tris-glycine-SDS running buffer and immunoblotting followed standard techniques using the following antibodies: mouse monoclonal anti-ISG15 F-9 (Santa Cruz Biotechnology Cat# sc166755), rabbit polyclonal anti-MxA (Proteintech Cat# 13750-1-AP), goat polyclonal anti-IFIT1 N-16 (Santa Cruz Biotechnology Cat# sc82946), mouse monoclonal anti-β-ACTIN, UBA7 (anti-UBE1L B-7; Santa Cruz Biotechnology Cat# sc-390097), rabbit monoclonal anti-phosphorylated STAT1 (anti-phospho-STAT1 (Tyr701) 58D6; Cell Signalling Technology Cat# 9167), mouse monoclonal anti-PIV5 NP 125 (24), mouse monoclonal anti-HPIV2 and anti-PIV5 P 161 (antibody cross-reacts with P of both viruses (24)), mouse monoclonal anti-HPIV3 NP (25). For quantitative immunoblots primary antibody-probed membranes were incubated with IRDye secondary antibodies (LiCOR) and signals detected using an Odyssey CLx scanner. Data were processed and analysed using Image Studio software (LiCOR).

### ^35^S-methionine labelling

Subconfluent A549 and A549-ISG15^-/-^ (B8) cells in 6-well plates were treated with 1000 IU/ml IFN-α or left untreated. At 24 h, 48 h and 72 h following treatment cells were pulse-labelled with 500 Ci/mmol ^35^S-Methionine (^35^S-Met; MP Biomedical) in Met-free media (Sigma) for 1 h. Cells were washed in PBS, lysed in 2 x Laemmli sample buffer and equal amounts of protein were separated by SDS-PAGE. Gels were stained with Coomassie (and imaged to ensure equal loading), dried under vacuum, exposed to a storage phosphor screen and analysed by phoshoimager analysis.

## Results

### ISG15-knockout A549 cells recapitulate ISG15-deficient patient cells

Among the several immune modulatory roles of ISG15 (8), intracellular ISG15 expression, at least in human cells, is critical for regulating the magnitude of the type I IFN response (10, 14). To investigate the pleotropic nature of human ISG15 we developed cell lines that lack ISG15 expression. Because of our interest in respiratory viruses, including paramyxoviruses, we chose to knockout ISG15 expression in the lung adenocarcinoma cell line A549 by CRISPR/Cas9 genome editing as described previously (18). Furthermore, A549 cells have proved to be a very useful model for understanding virus-IFN interactions. The resulting culture was single cell cloned and ISG15 expression was assessed by immunoblotting three clones (B8, B6 and C4). We also selected a clone that had gone through the CRISPR/Cas9 process but retained ISG15 expression (C4+) (Fig. 1a). In addition to control A549 cells, all clones were treated with IFN-α for 24 h, 48 h or left untreated. Immunoblot analysis showed that, compared to control cells, expression of the ISGs MxA and IFIT1 were higher in A549- ISG15^-/-^ cells (Fig. 1a). It was previously reported that increased ISG expression in ISG15-deficient cells was due to enhanced signalling resulting from the destabilisation of the type I IFN negative regulator USP18. To determine if IFN-α treatment led to enhanced signalling in A549-ISG15^-/-^ cells we selected clone B8 for further analyses. Cells were treated with IFN-α for 30 min, extensively washed and media without IFN-α was replaced. Immunoblot analysis of cell lysates taken after 30 min treatment (and following washes; 0’) and 30 min later (30’) showed that IFN-α treatment led to the phosphorylation of STAT1, an indicator of IFN signalling, in both A549 and A549-ISG15^-/-^ cells (Fig. 1b). Following 24 h treatment, there was clear evidence of ISG expression as shown by the expression of MxA and ISG15 (in A549 cells) and enhanced expression of STAT1 (Fig. 1b). However, while phospho-STAT1 levels had abated in both cell lines 24 h post-IFN-α treatment, levels were clearly higher in A549-ISG15^-/-^ cells indicating that in these cells there was a higher degree of signalling. We also tested the impact of ISG15-deficiency on the expression of various ISG mRNAs.

**Fig. 1.**
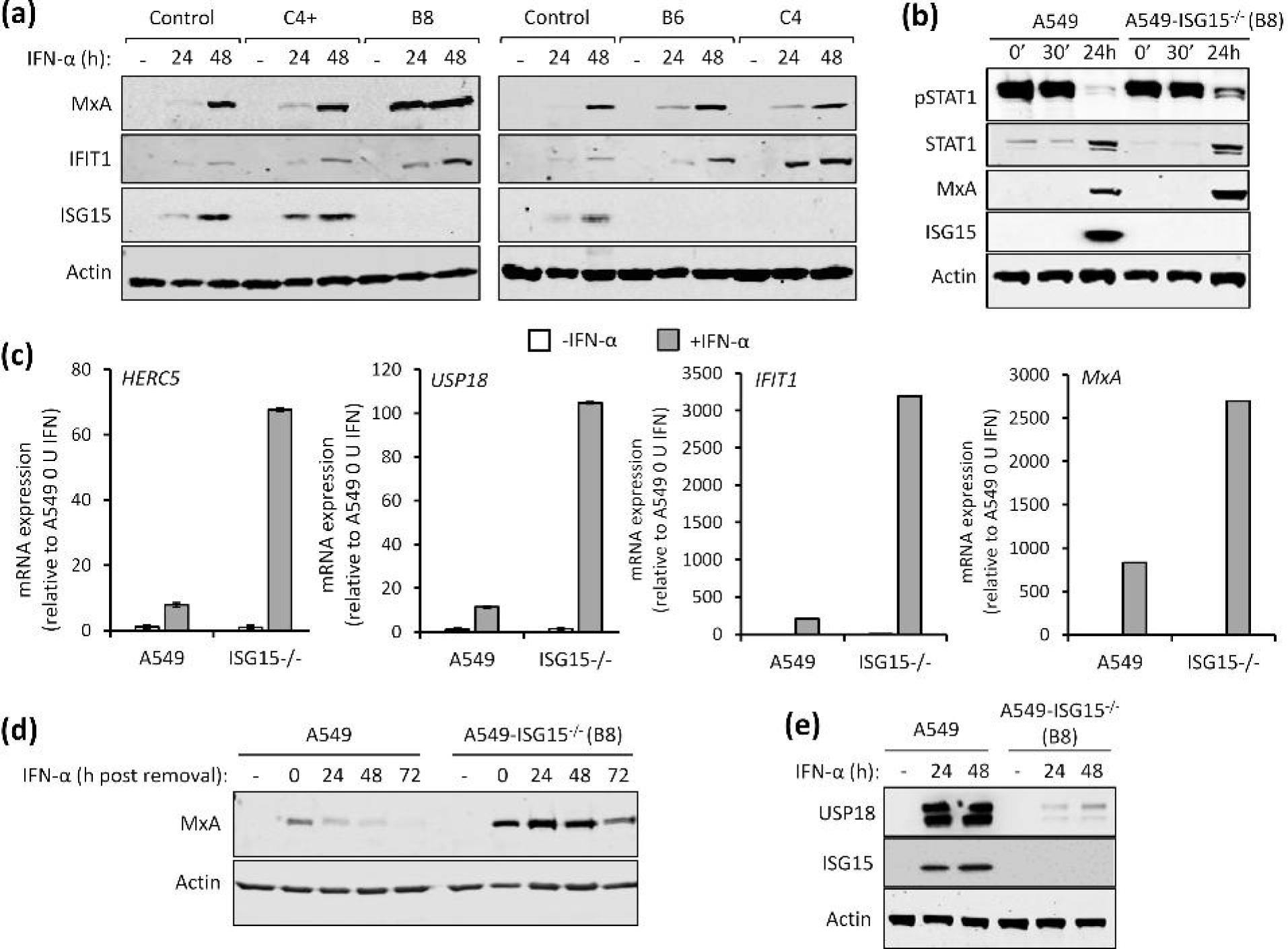
Functional characterisation of A549-ISG15 knockout cell lines. (a) CRSIPR/Cas9 genome editing was used to knockout ISG15 expression in A549 cells followed by single-cell cloning (following previously reported procedures (18)). Four independent clones were treated with 1000 IU/ml IFN-α for 24 and 48 h, or left untreated, and protein expression was tested by immunoblot analysis of ISG15, MxA, IFIT1 and β-Actin. ‘Control’ cells were naïve A549 cells. Representative image from two independent experiments. (b) A549 and A549-ISG15^-/-^ (B8) cells were treated with 1000 IU/ml IFN-α for 30 min then extensively washed and media without IFN replaced. Cells were harvested at 0 min (0’), 30 min (30’) and 24 h after IFN-α removal and phopho-STAT1, total STAT1, MxA, ISG15 and β- Actin were detected following immunoblot analysis. (c) A549 and A549-ISG15^-/-^ (clone B8) were treated with 1000 IU/ml IFN-α for 24 h. Expression of interferon stimulated genes was tested using RT-qPCR with primers specific for *HERC5, USP18, IFIT1* and *MxA*. Relative expression was determined following SYBR green qPCR using ΔΔCt method. β-Actin expression was used to normalise between samples. Error bars represent the standard deviation of the mean from three independent RNA samples. (d) A549 and A5549-ISG15^-/-^ (clone B8) were treated with 1000 IU/ml for 24 h. Cells were washed and fresh media (without IFN-α) was replaced. Cells were processed for immunoblot analysis using antibodies specific for MxA and β-Actin at 24 h post IFN-α and every 24 h thereafter for 72 h. Controls were cells without IFN-α. Representative image from two independent experiments. (e) A549 and A5549-ISG15^-/-^ (clone B8) were treated with 1000 IU/ml for 24 h and 48 h (or left untreated). Whole cell lysates were analysed by immunoblotting with antibodies specific for USP18, ISG15 and β-Actin. Image is representative of >3 independent experiments.

A549-ISG15^-/-^ cells were, in addition to control A549 cells, treated with IFN-α, or left untreated, for 24 h and the expression of various ISGs were examined by RT-qPCR. Whilst IFN-α treatment enhanced the expression of all ISGs tested, this increase was larger in ISG15-deficient cells compared to control A549 cells (between 5- and 10-fold, depending on the ISG) (Fig. 1c). Importantly, the expression of ISGs in non-stimulated cells was equivalent to control cells suggesting that ISG15- dependent regulation is specific to the IFN response and not required for the regulation of basal gene expression. Further experiments showed that lack of ISG15 prolonged the longevity of ISG protein expression, which presumably has an impact on patients with autoinflammatory diseases associated with ISG15 loss-of-function. Here, control A549 and knockout cells were treated with IFN- α for 24 h. The cells were washed and media (without IFN-α) was then added. Cells were harvested every 24 h for 72 h and MxA expression was assessed by immunoblotting (Fig. 1d). In control A549 cells MxA expression peaked at 24 h (the point at which IFN was removed) and had returned to basal levels between 48 and 72 h. In knockout cells MxA expression was clearly higher than in control cells, corroborating our mRNA analyses. Furthermore, while MxA expression in A549-ISG15^-/-^ did recede between 48 and 72 h, high protein levels remained at 72 h (Fig. 1d). A dysregulated IFN response in ISG15-deficient cells is thought to be due to destabilisation of USP18, a known negative regulator of JAK/STAT signalling (10). To determine if USP18 is similarly affected in our cell lines, A549-ISG15^-/-^ cells were treated with IFN-α for 24 or 48 h (or left untreated) and whole cell lysates were probed for USP18 by immunoblotting. USP18 was robustly induced in A549 cells following IFN-α treatment; however, levels of USP18 were much lower in IFN-α-treated ISG15-deficient cells (Fig. 1e). *USP18* mRNA levels were approximately 10-fold higher in IFN-treated ISG15-deficient cells compared to control A549s, demonstrating that reduced USP18 in A549-ISG15^-/-^ cells was not due to reduced transcription (Fig. 1c). Together, these data show that ISG15 is critical for the regulated expression of ISGs. Moreover, they demonstrate that the effects of IFN treatment on our ISG15 knockout A549 cell lines recapitulate the findings in cells derived from ISG15-deficent, patient cells.

### ISG15-deficiency leads to translational repression following IFN treatment

During our studies we observed that IFN-α-treatment of ISG15-knockout cells led to a reduction in protein synthesis and reasoned that this was a likely contributor to the reported virus resistance in ISG15-deficient cells (14). To investigate this we treated, or left untreated, A549 and A549-ISG15^-/-^ (B8) cells with IFN-α. At 24 h, 48 h and 72 h following treatment cells were pulse labelled with ^35^S- Methionine (^35^S-Met) for 1 h and the incorporation of ^35^S-Met was analysed by phoshoimager analysis. These data showed, compared to control cells, that there was a pronounced decrease in protein synthesis in ISG15^-/-^ cells between 24 h and 48 h (Fig. 2a). We also investigated whether this decrease in protein synthesis would lead to the inhibition of viral protein synthesis. Cells were pre-treated with IFN-α for 8 h, or left untreated, infected with the orthorubulavirus PIV5 (family *Paramyxoviridae*, sub-family *Orthorubulavirinae*) at a MOI of 10 and then labelled for 1 h with ^35^S- Met at 24 h and 48 h p.i. (32 h and 56 h post IFN-α treatment, respectively). Because of the abundance of viral proteins in infected cells, they can be observed by phophorimager analysis, which, following a 1 h treatment of infected cells with ^35^S-Met at 24 and 48 h p.i., showed higher levels of newly synthesised viral protein at 24 h p.i. than at 48 h p.i. in A549 cells (Fig. 2b). This is because peak viral transcription occurs between 18 and 24 h p.i. (26). This differs from immunoblot analysis that measures the accumulation of viral protein over time; here, the levels of viral protein appeared as high, if not higher, at 48 h p.i. than 24 h p.i. (Fig. 2c). In contrast, the levels of viral protein synthesis following IFN-α treatment was higher at 48 h p.i. than at 24 h p.i. because IFN-α treatment delayed PIV5 infection (Fig. 2b). This was also indicated by immunoblot analysis where the accumulation of NP was higher at 48 h p.i. that 24 h p.i. (Fig. 2c). When A549-ISG15^-/-^ cells were infected, there was clear evidence of NP protein synthesis (Fig. 2b) and accumulation (Fig. 2c); however, when these cells were pre-treated with IFN-α and infected, there was very little evidence of viral protein synthesis (Fig. 2b) or accumulation (indicating that viral protein synthesis was barely initiated) (Fig. 2c) at any time p.i. These data demonstrate that IFN-α-treatment of A549-ISG15^-/-^ cells led to inhibition of protein synthesis that was associated with viral resistance, at least at later times.

**Fig. 2.**
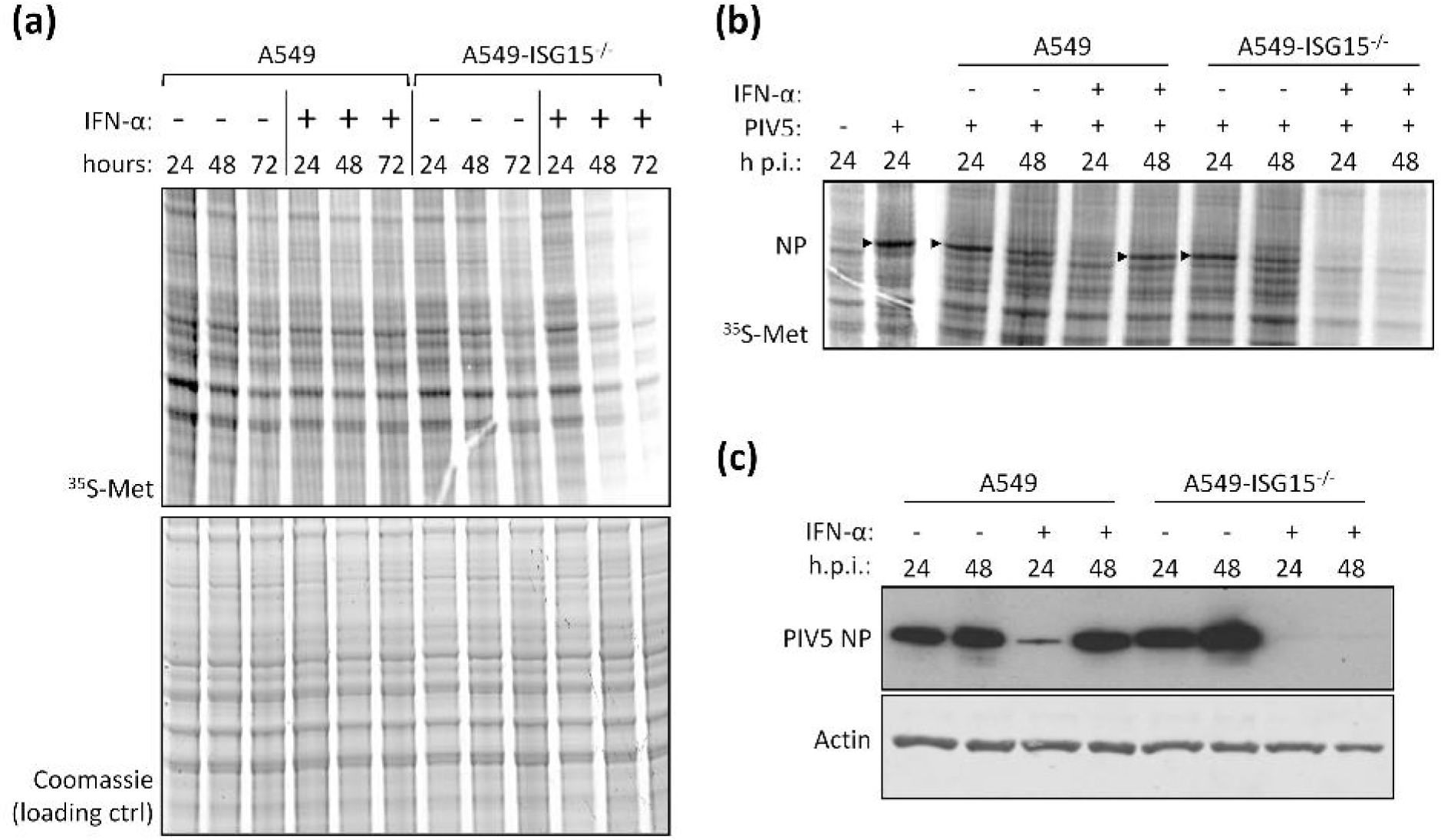
Analysis of cellular and viral protein synthesis in ISG15-deficient cells during an antiviral state. (a) Sub-confluent A549 and A549-ISG15^-/-^ (B8) cells were treated with 1000 IU/ml IFN-α or left untreated. At 24 h, 48 h and 72 h cells were pulsed for 1 h with L-[^35^S]-Methionine (^35^S-Met) in Met-free media to metabolically label nascent proteins. Proteins were resolved by SDS-PAGE and stained with Coomassie to ensure equal loading. Labelled proteins were visualised by phophoimager analysis. (b) A549 and A549-ISG15^-/-^ (B8) cells were treated with 1000 IU/ml IFN-α for 8 h or left untreated and then infected with PIV5 strain W3 (MOI = 10). At 24 or 48 h post infection cells pulsed and processed as in (a). Arrow heads denote ^35^S-Met-labelled PIV5 nucleoprotein (NP). Both experiments were performed independently at least twice. (c) PIV5-infected lysates from (b) were immunoblotted and the accumulation of PIV5 NP and β-Actin were detected with specific antibodies and HRP-conjugated secondary antibodies.

### Pre-treatment of ISG15-deficient cells with IFN-α renders them resistant to parainfluenza virus infection

Previous studies have shown that IFN-α treatment of ISG15-deficient patient cells renders them resistant to viral infection by several viruses, including the murine respirovirus (family *Paramyxoviridae*, sub-family *Orthoparamyxovirinae*) Sendai virus (14), and this seems to extend to PIV5 with our *in vitro* system (Fig. 2b-c). To investigate this in A549-ISG15^-/-^ cells, control A549 cells and the ISG15 knockout clones described above were either untreated or treated with 1000 IU/ml IFN-α2b (the same concentration and IFN-α type used in (14)) for 18 h. Cells were then infected with PIV5 (strain W3) (22) for 24 and 48 h and analysed by immunoblotting. In all cell lines, the levels of PIV5 nucleoprotein (NP) expression was equivalent at 24 and 48 h in unstimulated cells (Fig. 3a). In IFN-α pre-treated control cells, including C4+ that retained ISG15 expression, the level of NP expression was markedly reduced at 24 h. By 48 h, the level of NP increased showing that infection had progressed even in the presence of IFN-α (Fig. 3a). This is because the PIV5-V protein targets STAT1 for proteasomal degradation, and once sufficient V is expressed, the IFN response is dismantled allowing the virus to replicate (23). Indeed, there was no detectable STAT1, and as a result, markedly reduced levels of ISGs MxA and IFIT1 in PIV5-infected, ISG15-expressing cells (Fig. 3a). However, all A549-ISG15^-/-^ cell lines that had been pre-treated with IFN-α were resistant to PIV5 infection as shown by dramatically reduced, or even absent, NP expression at both time points (Fig. 3a). Moreover, these cells displayed STAT1 expression and the expression of associated MxA and IFIT1 (indicating that PIV5 infection was inhibited) (Fig. 3a).

**Fig. 3.**
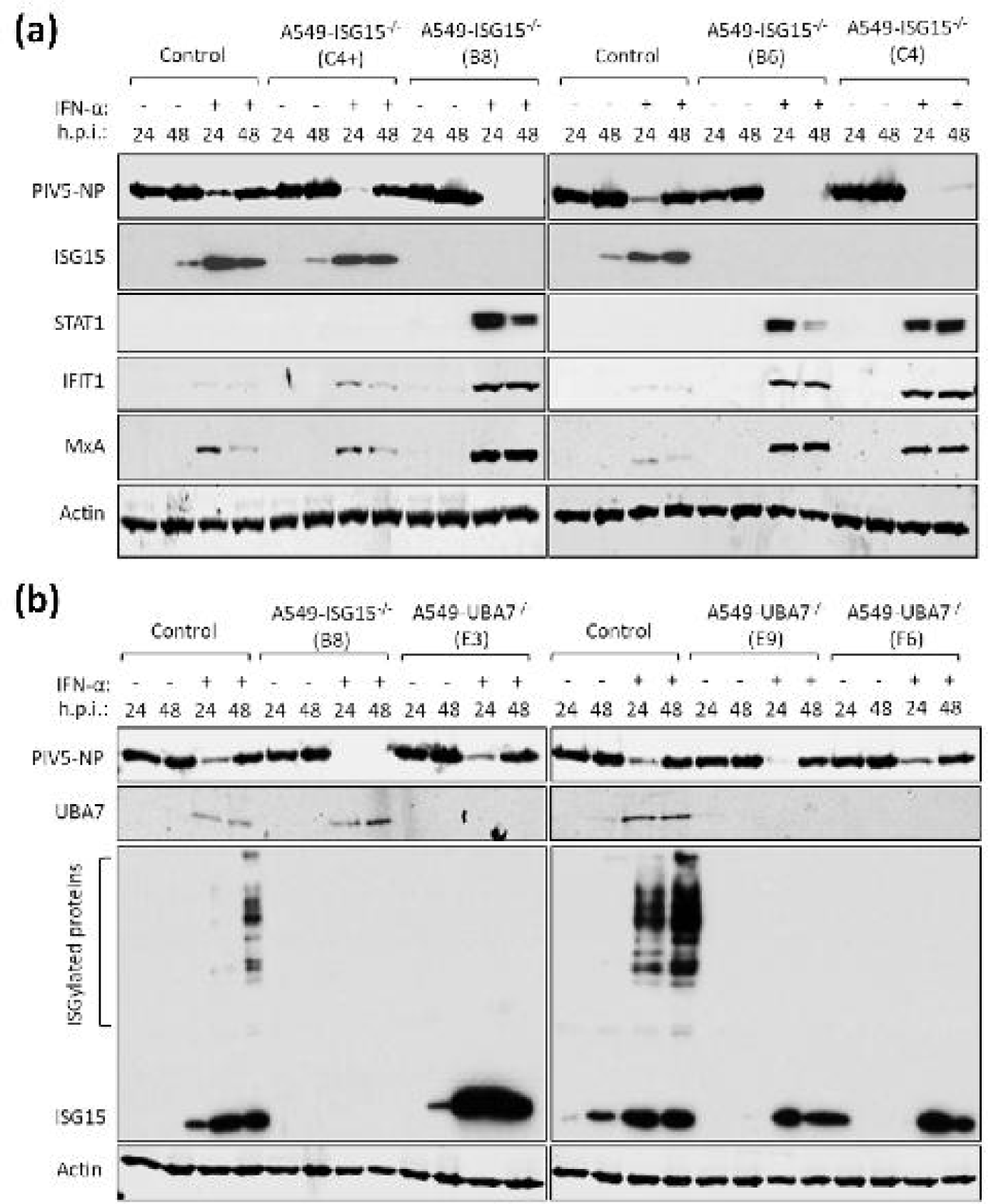
IFN-α pre-treatment of ISG15-deficient cells leads to virus resistance which is independent of ISGylation. (a) Control (naïve A549) and 4 independent clones of A549-ISG15^-/-^ cells generated by CRISPR/Cas9 genome editing were treated with 1000 IU/ml IFN-α for 16 h or left untreated and then infected with PIV5 strain W3 (MOI = 10). Cells were harvested at 24 h and 48 h p.i. and processed for immunoblot analysis using antibodies specific for PIV5 nucleoprotein (NP), ISG15, STAT1, IFIT1, MxA and β-Actin. This experiment was independently performed twice. (b) UBA7 knockout cells were generated using CRISPR/Cas9 genome editing; Cas9-expressing A549 cells were first generated (following transduction with lentiCas9-Blast) and then transduced with lentiGuide-Puro expressing a single guide RNA that targeted exon 3 of the UBA7 gene. Knockout cells were single cell cloned and three were selected for further analysis. These cells were treated with IFN-α or left untreated, infected and processed as in (a) using antibodies specific for PIV5 NP, ISG15, UBA7 and β-Actin. This experiment was independently performed twice.

Previous reports have shown that the ISG15 regulation of IFN signalling is independent of its ability to covalently modify proteins by ISGylation (10). To confirm this, we again applied CRISPR/Cas9 genome engineering technology and knocked out expression of UBA7, the E1 enzyme required for ISGylation. For this we took a different approach compared to generating our ISG15 knockout cells (19). Here, we introduced constitutive expression of Cas9 by lentiviral transduction of A549 cells and transduced A549-Cas9 cells with lentiGuide-Puro lentivirus carrying a guide RNA specific for UBA7, followed by single-cell cloning. We confirmed that all clones were UBA7-deficient by immunoblot analysis, which demonstrated that they retained expression of ISG15 but had lost the ability to ISGylate proteins (Fig. 3b). Additionally, following the scheme used in Fig. 3a, these cells were infected with PIV5-W3. These data showed that, compared to ISG15 knockout cells that were resistant to infection, all IFN-α-pre-treated UBA7-knockout cells were infected as efficiently as control cells (Fig. 3b), confirming reports that ISG15-dependent regulation of type I IFN signalling does not require ISGylation (10).

### The direct antiviral activity of ISGs is responsible for virus resistance

Virus resistance can be induced following 8 h IFN-α treatment (shorter times were not tested), well before any obvious effect on global protein synthesis (Fig. 2). Therefore, shutdown of translation is unlikely to be the sole contributor to virus resistance at early time points and so we wished to determine whether the direct antiviral activity of ISGs was responsible. Addressing this question is complex since, for most viruses, the specific ISG(s) responsible for blocking replication is not known. However, for PIV5, it has been established that IFIT1 is the principle ISG responsible for most of the IFN-dependent antiviral activity (17, 27). We therefore hypothesised that if virus resistance was caused by the direct antiviral activity of ISGs, knockdown of IFIT1 in ISG15-deficient cells would permit PIV5 replication during an antiviral response. We reduced IFIT1 (according to (17)) in A549 and A549-ISG15^-/-^ cells and all four cell lines (A549, A549-ISG15^-/-^ and the respective shIFIT1 cells) were pre-treated, or left untreated, with IFN-α and then infected with PIV5-W3 (MOI 10) for 24 and 48 h. Expression of PIV5 NP, analysed by semi-quantitative immunoblotting, was used to measure virus infection (Fig. 4a). IFIT1 levels and expression of ISG15 were likewise tested. Typically, pre-treatment of naïve cells with IFN-α reduced infection, as shown by a reduction in NP levels, compared to non-treated cells (Fig. 2b-c & 3a-b); nevertheless, because PIV5 expresses the IFN antagonist V protein, NP levels reach similar levels to untreated cells by 48 h p.i. However, this IFN-dependent reduction in virus infection is diminished when IFIT1 is knocked down, confirming earlier reports of IFIT1’s antiviral activity against PIV5 (17, 27). While IFN-α pre-treatment of A549-ISG15^-/-^ cells renders them resistant to infection, when IFIT1 was also knocked down, PIV5 infection was restored (Fig. 4a). Because we performed semi-quantitative immunoblotting of NP and β-Actin, we were able to quantify NP levels, allowing us to analyse these changes statistically (Fig. 4b). These data show that in IFN-α-pre-treated cells, knocking IFIT1 down restored NP to similar levels to those seen in untreated cells, regardless of ISG15 status. While IFN-α pre-treatment of A549 cells significantly reduced NP levels when we compared 24 h and 48 h p.i. samples, there was no difference at these time points when IFIT1 was knocked down (Fig. 4b). Importantly, while NP levels were virtually absent in IFN-α-pre-treated ISG15-deficient cells, when IFIT1 was knocked down in these cells NP levels were equivalent to A549-shIFIT1 cells (Fig. 4b).

**Fig. 4.**
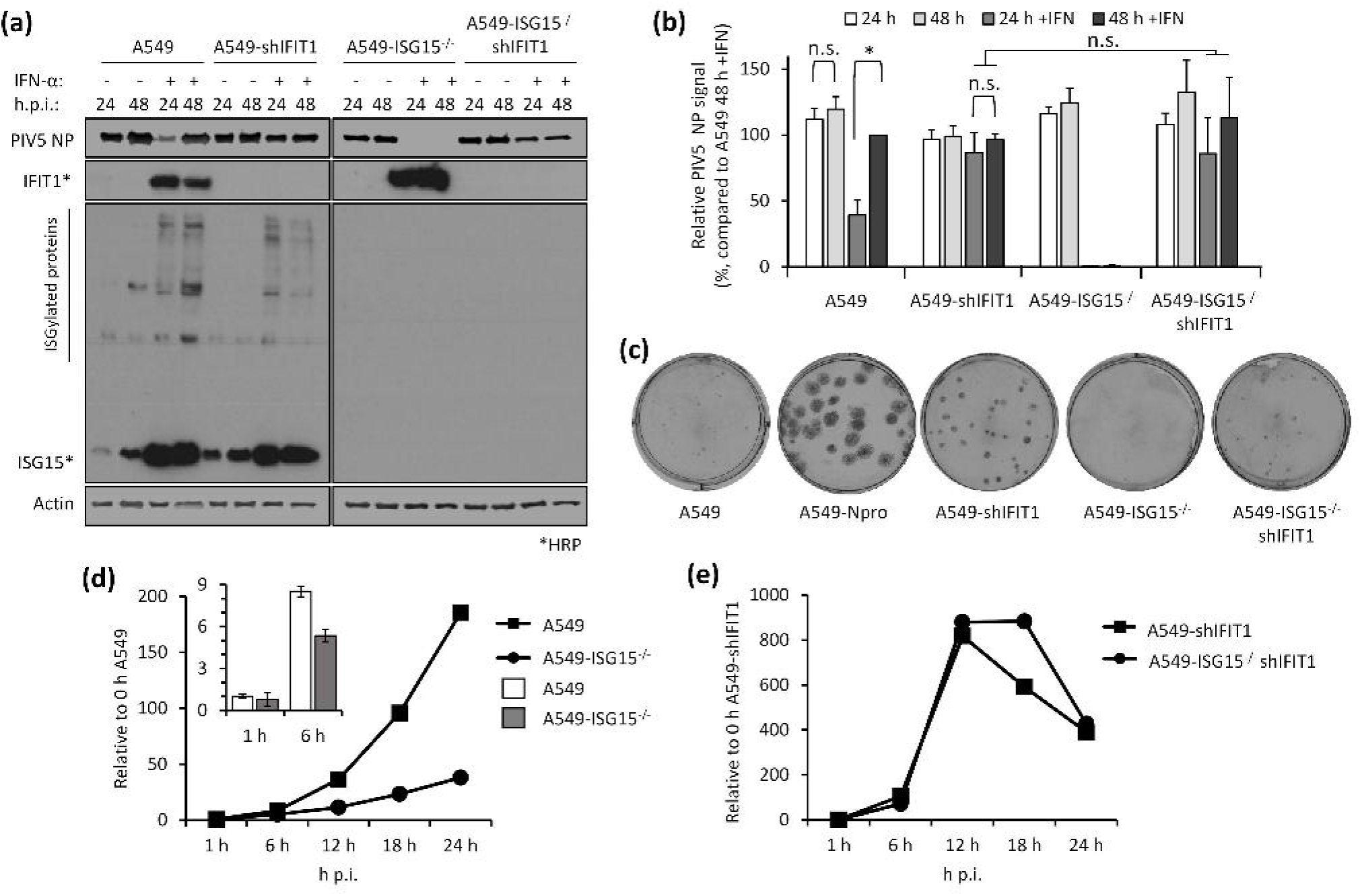
Direct antiviral activity of ISGs is responsible for virus resistance due to ISG15-loss-of-function. (a) IFIT1 was constitutively knocked down in A549 or A549-ISG15^-/-^ (B8) cells following a previously described method (17). A549, A549-ISG15^-/-^ (B8) and the corresponding IFIT1 knockdown cells were treated with IFN-α, infected with PIV5 and processed as in Fig. 3a. Following immunoblotting with specific antibodies, PIV5 NP and β-Actin were detected using near-infrared (NIR) dye-conjugated secondary antibodies to facilitate quantification. IFIT1 and ISG15 proteins were detected using chemiluminescence following incubation with horseradish peroxidase (HRP)-conjugated secondary antibodies. (b) Experiments described in (a) were performed independently three times (infections were performed on three separate occasions) and NP and β-Actin levels were quantified using Image Studio software (LiCOR). Signals were relative to those generated from IFN-α-treated A549 cells infected for 48 h p.i. (set to 100%). Error bars represent the standard deviation of the mean from the three independent experiments performed on different occasions. Asterisks denote statistical significance using two-way ANOVA and Tukey’s multiple comparisons test: * (P < 0.05), n.s. denotes no statistical significance. (c) Indicated cells were infected for 1 h with 30 – 40 plaque forming units (PFU) of PIV5 (CPI-), a strain unable to block the IFN response due to mutation in the viral V protein. Monolayers were fixed 6 d p.i. Plaques were detected using a pool of anti-PIV5 antibodies specific for hemagglutinin (HN), nucleoprotein (NP), phosphoprotein (P) and matrix protein (M) (see (24)). Plaque assays were performed on 3 independent occasions. (d) A549 and A549-ISG15^-/-^ cells were infected with PIV5 W3 (MOI 10) and harvested at the indicated times. Total RNA was isolated and subjected to cDNA synthesis using oligo(dT) primers. Expression of PIV5 NP was measured using qPCR. Relative expression (compared to 1 h A549) was determined following SYBR green qPCR using ΔΔCt method. β-Actin expression was used to normalise between samples. Error bars represent the standard deviation of the mean from three independent RNA samples. For clarity, the inset bar graph represents viral transcription data at 1 h and 6 h p.i. only. (e) Analyses followed that of (d) but A549-shIFIT1 and A549-ISG15^-/-^/shIFIT1 cells were infected.

Rather than solely relying on viral protein expression as a surrogate for virus infection, we also tested virus replication using biologically relevant plaque assays. Because paramyxoviruses (like most wild type viruses) are poor inducers of the IFN response (28, 29), are able to efficiently and rapidly counteract it if it were induced, and our data showed that basal ISG expression was not effected in ISG15-deficient cells (Fig. 1c), we predicted that infection of naïve A549-ISG15^-/-^ cells would be equivalent to naïve A549 cells. To determine if this was the case, plaque assays were performed with various paramyxoviruses. These data show that each virus formed plaques that were analogous on both A549 and A549-ISG15^-/-^ cells (Supplemental Fig. 1). There were subtle differences in plaque phenotype; for instance, infection of ISG15-deficient cells, particularly with HPIV2 but also evident following PIV5 infection, resulted in plaques with poorer defined edges (hazy plaques) (Supplemental Fig. 1). The reason for this is currently not clear but may indicate an antiviral role for ISG15 against HPIV2 and PIV5. Nevertheless, this, and data in figures 2 and 3, supports the notion that naïve cells were not resistant to wild type viral infection. However, viruses unable to counteract the IFN response should be restricted and therefore provide a means of assessing the role of ISG15 and virus resistance.

To do this cells were infected with approximately 30-40 PFU of PIV5 strain CPI- (PIV5-CPI-) (30), a strain unable to block IFN signalling due to a mutation in its V protein. Infected cells were fixed 6 d p.i. and stained for viral antigen (Fig. 4c). As previously demonstrated (17), PIV5-CPI- was unable to efficiently form plaques in IFN-competent A549 cells. However, PIV5-CPI- did replicate when cells were unable to produce IFN, such as in A549-Npro cells that constitutively express bovine viral diarrhea virus (BVDV) Npro that cleaves IRF3 (a transcription factor critical for IFN induction (21)). Furthermore, when IFIT1 was knocked down, PIV5-CPI- was able to replicate (albeit less efficiently), further highlighting the major role of IFIT1 as an anti-PIV5 protein. As expected, and like A549 cells, there was very little virus replication in A549-ISG15^-/-^ cells; however, when IFIT1 was knocked down, cells were able to support virus replication. It must be noted however that virus replication in A549- ISG15^-/-^/shIFIT1 cells did not recover to the same degree as A549-shIFIT1 cells. We propose that the reason for this will be complex and may include the likelihood that additional, yet to be identified, anti-PIV5 ISGs exist which are expressed at higher levels in ISG15-deficient cells. Another possible explanation is the inhibition of protein synthesis, including that of viral proteins, in ISG15-deficent cells; cells were infected for 6 days prior to performing the plaque assays, a time point beyond that required to observe a significant effect on protein synthesis (Fig. 2a). Therefore, the plaques observed in A549-ISG15^-/-^/shIFIT1 cells likely result from virus that replicated prior to the inhibition of global protein synthesis.

IFIT1 restricts viral infection post-transcriptionally by blocking the translation of viral mRNA (17, 27); therefore, we predicted that IFN-α-pre-treated A549-ISG15^-/-^ cells would remain susceptible to infection, but that high levels of IFIT1 would mean these cells would not be permissive to PIV5 infection. Furthermore, investigating this could highlight additional restrictions to viral infection, such as entry. A549 and A549-ISG15^-/-^ cells were pre-treated for 8 h with IFN-α and then infected with PIV5-W3 (MOI 10) (Fig. 4d). Analysis of PIV5 *NP* transcription showed that ISG15-deficent cells were infected and that viral transcription increased over time; however, this was muted compared to A549 control cells. Importantly however, the levels of *NP* transcription at 1 h p.i. was equivalent in both cell lines, a time point that likely represents primary transcription (Fig. 4d; see inset graph). These data suggest that both cell lines were susceptible to infection and that high levels of pre-existing IFIT1 strongly restricted further viral transcription by preventing the translation of the virally encoded mRNAs. To investigate if IFIT1 restriction was responsible for reduced viral transcription in ISG15-deficient cells, we repeated the experiment in A549-shIFIT1 and A549-ISG15^-/-^/shIFIT1 cells (Fig. 4e). These data show that in IFN-α-treated cells, viral transcription was markedly increased compared to cells with intact IFIT1 expression. Furthermore, in A549-shIFIT1 cells, transcription peaked between 12 and 18 h p.i. and then receded. We have recently described the transcription and replication of various paramyxoviruses, including PIV5-W3, using un-biased high throughput, RNA-seq approach (26); this report shows that this pattern of transcription is typical of PIV5-W3 and likely results from the phosphoprotein (P)-dependent repression of viral transcription and replication (31). This repression also occurred in A549-ISG15^-/-^/shIFIT1 cells, but this occurred later (Fig. 4e), suggesting that ISG15 may be an additional antiviral factor that curtail PIV5 transcription. Nevertheless, these data showed that when IFIT1 levels were knocked down, the transcriptional repression identified in IFN-α-pre-treated ISG15-deficient cells was relieved, demonstrating that virus resistance was due to the post-transcriptional activity of IFN-inducible IFIT1. We also investigated infection of these cell lines with other paramyxoviruses whose sensitivity to IFIT1 has been previously reported. Cells were treated with IFN-α and then infected with HPIV2 strain Colindale (MOI 10; family *Paramyxoviridae*, sub-family *Orthorubulavirinae*), which is reported to be moderately sensitive to IFIT1-restriction (27), for 24 and 48 h (untreated cells were not analysed because of high cytopathic effect in the absence of IFN). To investigate infection, we detected expression of HPIV2 phosphoprotein (P) by semi-quantitative immunoblotting (Fig. 5a), which showed that IFN-α-pre-treated A549-ISG15^-/-^ cells were largely resistant to infection, although by 48 h p.i. there was some, albeit low level, evidence of viral protein accumulation. Nevertheless, infection of A549-ISG15^-/-^/shIFIT1 did allow significantly more viral protein expression. Semi-quantitative analyses demonstrated that viral protein accumulation in A549-ISG15^-/-^/shIFIT1 cells was significantly higher than in A549-ISG15^-/-^ cells, but this was not as high as in A549 control cells, which agrees with the reported partial sensitivity of HPIV2 to IFIT1 restriction indicating that additional ISGs target HPIV2 (Fig. 5b). We performed a similar analysis with HPIV3 strain Washington (20) (family *Paramyxoviridae*, sub-family *Orthoparamyxovirinae*), a virus reported to have limited sensitivity to IFIT1 (27). Interestingly, pre-treatment of A549 and A549-shIFIT1 cells with IFN-α had less of an effect on virus protein accumulation compared to the effects on PIV5 infection (Fig. 5c). Furthermore, while infection of IFN-α-pre-treated ISG15 knockout cells significantly reduced infection compared to control cells, virus infection in these cells was still more robust compared to PIV5 and HPIV2-infected cells. Nevertheless, knockdown of IFIT1 only slightly increased HPIV3 protein expression in both ISG15-competent and ISG15-deficient cells (Fig. 5d), supporting reports of a minor role of IFIT1 during the antiviral response to HPIV3 (27).

**Fig. 5.**
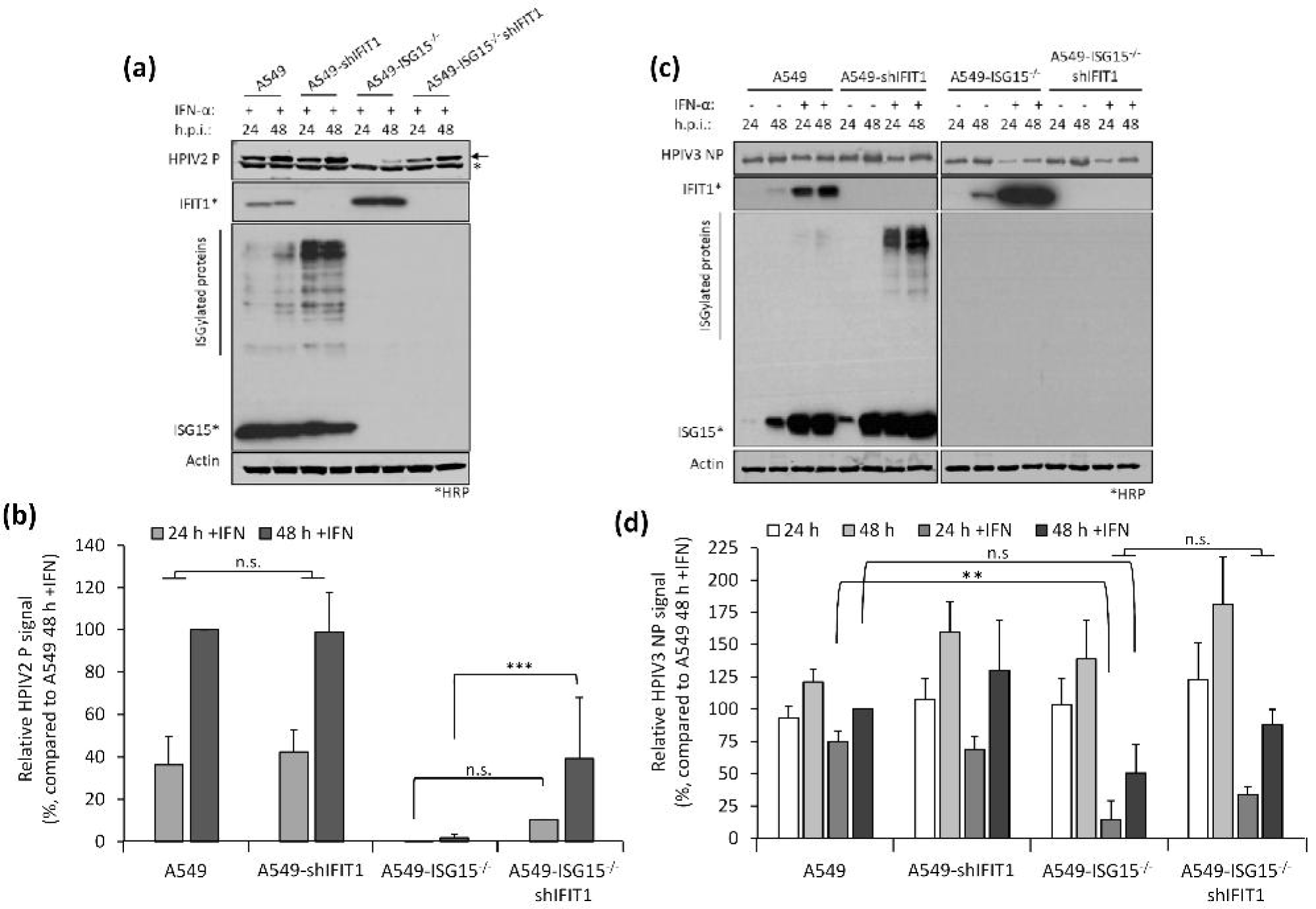
Restoration of paramyxovirus infection in IFN-α-pretreated ISG15^-/-^ cells reflects their reported sensitivity to IFIT1. (a-b) Experiments were performed as in (Fig. 4a-b). (a) HPIV2 proteins were detected with antibodies specific for HPIV2 P (clone 161; (24)) and NIR-conjugated secondary antibodies. Asterisks denotes detection of an irrelevant protein, arrow denotes HPIV2 P. Samples not treated with IFN-α were omitted due to the highly lytic nature of HPIV2 which hampered their accurate quantification. (b) Quantification of normalised NP signals and compared to the 48 h p.i. sample that was set to 100%. (c) HPIV3 NP proteins were detected using antibodies specific for HPIV3 NP and NIR-conjugated secondary antibodies. (d) Normalised signals were quantified as in (b) and compared to IFN-α-treated, 48 h p.i. samples (set to 100%). Means and standard deviations were derived from 5 independent experiments for HPIV2 and 4 independent experiments (for HPIV3) performed on different occasions. Asterisks denote statistical significance using two-way ANOVA with Tuckey’s multiple comparison test (for HPIV3) and one-way ANOVA with Sidak’s multiple comparisons test (for HPIV2): ** (P < 0.01), *** (P < 0.001), n.s. denotes no statistical significance.

### ISG15-deficient cells pre-treated with IFN-α for longer times were resistant to infection independently of the direct antiviral activity of IFN-dependent restriction factors

Our data have so far suggested that early virus resistance is mediated by the direct antiviral activity of the IFN response. However, protein synthesis is reduced at later times post-IFN treatment and this is likely to cause resistance; therefore, we investigated whether PIV5 resistance could be induced independently of the direct antiviral activity of IFIT1. To do this we pre-treated the four cell lines (A549, A549-shIFIT1, A549-ISG15^-/-^ and A549-ISG15^-/-^-shIFIT1) with IFN-α for different periods of time, infected with a recombinant PIV5 that expresses the fluorescent protein mCherry (rPIV5-mCherry) for 48 h (MOI 10) and measured fluorescence as a marker of virus replication (Fig. 6a). Virus replication in A549 cells was equivalent regardless of the time cells had been pre-treated with IFN-α and, as expected, A549-ISG15^-/-^ cells were resistant to infection at any time post IFN-α treatment (Fig. 6b). Any advantage to PIV5 replication as a result of IFIT1 knockdown in A549-shIFIT1 cells was lost when cells had been pre-treated for 16 h or more, as longer periods of pre-treatment resulted in replication equivalent to IFN-pre-treated A549 cells. Similarly, PIV5 replication in A549- ISG15^-/-^-shIFIT1 cells was higher than A549 control cells, and equivalent to A549-shIFIT1 cells, following 8 and 16 h pre-treatment; however, when cells were pre-treated for 24 h, replication was lower than in A549 and A549-shIFIT1 cells. Interestingly, as the time of pre-treatment of A549-ISG15^-/-^-shIFIT1 cells extended, virus replication reduced further until cells became resistant (e.g. at 60 h and 72 h pre-treatment, Fig. 6b), which was not observed in A549 or A549-shIFIT1 cells. These data suggest that cell permissiveness progressively reduced with longer times of IFN-α pre-treatment, which correlated with the effects of IFN-α treatment on protein synthesis in ISG15-deficient cells (Fig. 2).

**Fig. 6.**
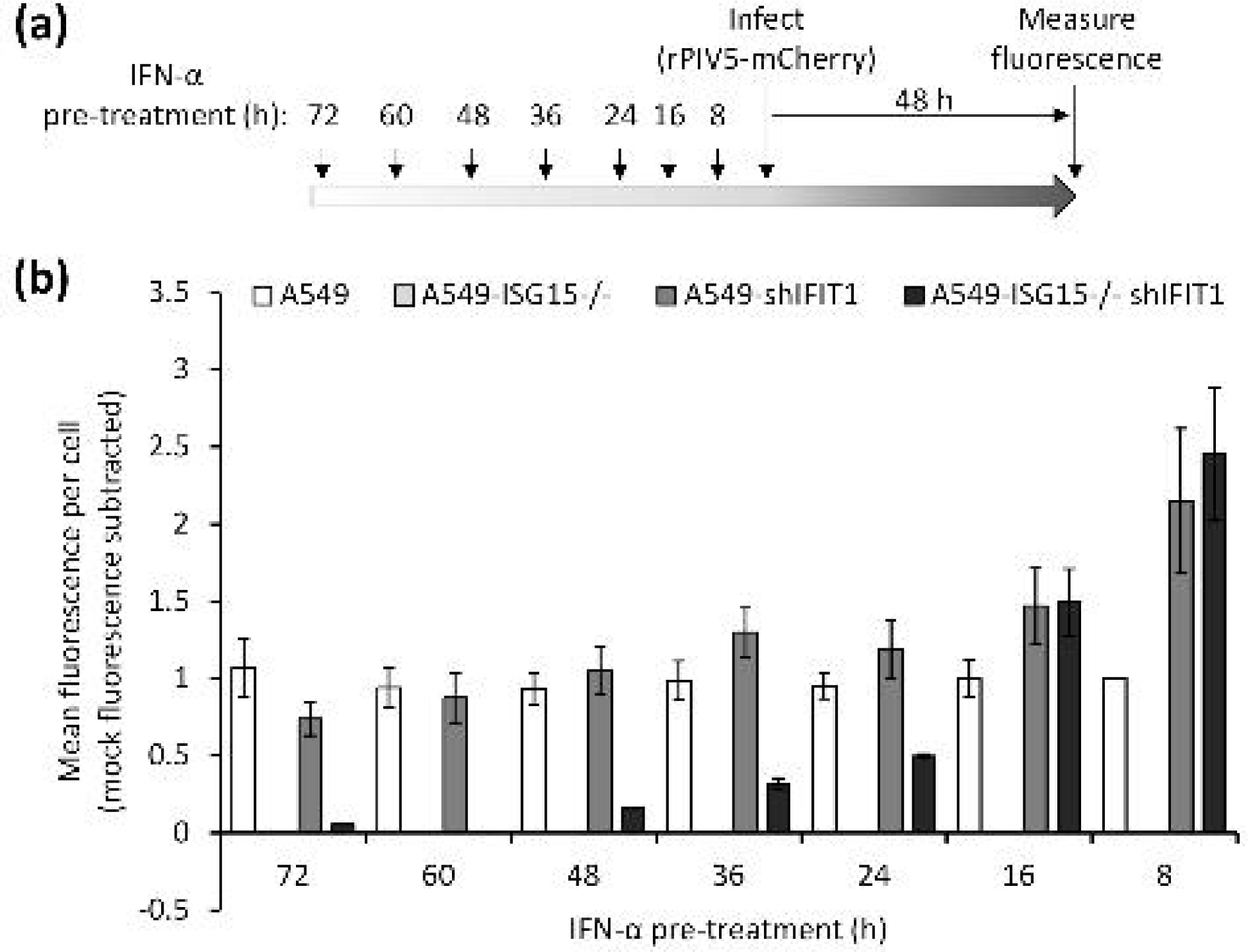
Virus resistance is induced in ISG15-deficient cells following longer periods of IFN-α pre-treatment. (a) Experimental workflow. (b) Cells were treated 1000 IU/ml IFN-α in 6-well plates for the indicated times prior to infection. Pre-treated cells were infected with rPIV5-mCherry (MOI 10) for 48 h and mCherry fluorescence was measured using an IncuCyte ZOOM. Background fluorescence from mock-infected wells was subtracted. Data are representative to two independent experiments.

A previous report demonstrated that ISG15-dependent stabilisation of USP18 was required to bring about regulation of the type I IFN response and this was sufficient for these cells to once again be infected (14). However, what aspects of the antiviral response was responsible for resistance was not investigated. Taken together, these data strongly suggest that virus resistance in early IFN-treated ISG15-deficient cells was caused by the direct antiviral activity of ISGs and not due to a lack of permissiveness as a result of IFN-dependent inhibition of protein synthesis. Nevertheless, because of the reduced protein synthesis in IFN-α-treated ISG15-deficient cells, cells later become non-permissive to infection, even when key ISGs are eliminated.

## Discussion

Previous work had shown that virus resistance was observed in cells that had been treated with IFN- α and then left to rest for 36 h prior to challenge (14). We had observed that IFN-α treatment of A549-ISG15^-/-^ cells led to dramatic decreases in protein synthesis, particularly between 24 and 48 h; therefore, it was not clear whether the initially reported virus resistance was due to defects in translation (including of viral mRNAs) at the timepoint used in (14) or due to the direct antiviral activity of the IFN response. For most viruses, the specific ISG(s) with antiviral activity for a given virus is not known, making the latter difficult to discern; however, for PIV5, it is well established that IFIT1 is responsible for the majority of the antiviral response (17). To study this we generated A549- ISG15^-/-^ cells and showed these cells recapitulated the effects observed in ISG15-deficient patient cells following treatment with IFN-α which included dysregulated ISG expression and reduced USP18 protein levels following IFN-α treatment (Fig. 1). Additionally, by knocking-out UBA7, the first enzyme in the ISGylation cascade, we showed that ISGylation is not required for a regulated response (Fig. 3b), confirming previous reports that ‘free’ ISG15 is required for regulation (10).

Using these cell lines in combination with a PIV5 infection model, we showed that infection of IFN-α- pre-treated ISG15-deficient cells in which IFIT1 had been knocked down restored infection, thus confirming that at early times post infection, resistance was indeed due to the direct antiviral activity of the IFN response. Furthermore, because IFIT1 blocks the translation of viral transcripts, our data show that IFN-treated A549-ISG15^-/-^ cells were still susceptible to infection, allowing viral transcription to take place prior to IFIT1 restriction, and that ISG15 was unlikely to significantly regulate processes involved in entry (Fig. 4d-e). Nevertheless, if ISG15-deficient cells were treated for longer periods with IFN-α prior to infection they did become resistant, even when IFIT1 was knocked down, suggesting that at later times the inhibition of protein synthesis was the principal cause of resistance (Fig. 6). These data suggest that the virus resistance reported by Speer et al. (14) was due to a lack of permissiveness and not a result of the direct antiviral activity of the IFN response, although different cells were used in that study.

The data here demonstrate that the mechanism of resistance is likely two-fold, depending on the duration that cells are exposed to IFN-α. It is not currently possible to know which mechanism is dominant in ISG15-deicient patients, but it is likely to be a combination of both. Nevertheless, virus resistance results from a lack of IFN signalling control - as a consequence of ISG15-loss-of-function - which would explain why ISG15-deficient patients were not more susceptible to severe infection. This observation, therefore, cannot be used to support the notion that human ISG15 does not possess direct antiviral activity, as proposed (14, 16). It is likely that many viruses will not be sensitive to ISG15-dependent antiviral activity; however, this is true of many antiviral effectors. For example, and as confirmed in this study, IFIT1 strongly restricts PIV5 infection, yet it has reduced activity against HPIV2 and likely no activity against HPIV3 or human respiratory syncytial virus (27). It is also true that several ISGs are often required to limit infection (6); therefore, if one antiviral effector mechanism is absent (such as ISGylation), there is sufficient redundancy to avoid severe effects of infection (redundancy that can complicate the investigation of specific antiviral mechanisms in *in vitro* studies). Nevertheless, several human viruses have been shown to be sensitive to ISGylation and many have evolved specific mechanisms to counteract antiviral ISGylation, adding further weight to the argument that human ISG15 does have antiviral activity (reviewed in (8)). Indeed, other than the handful of patients that have been found to lack ISG15 expression (10, 32), individuals will possess an intact IFN response where the antiviral activity of ISG15 (and other effectors) will function, if the infecting virus is sensitive to it.

It was surprising that protein synthesis was so affected in ISG15-deficient cells following IFN treatment. It is well established that inhibition of general protein translation is a key feature of the antiviral response and this is through the actions of proteins such as PKR or PERK (PKR-like ER kinase) (4). However, for PKR to be activated it must recognise dsRNA, which was absent in IFN-α-treated cells. Similarly, PERK is activated upon endoplasmic reticulum stress which might be expected during a viral infection, but not following treatment with IFN alone. Previous reports have shown that carcinoembryonic antigen-related cell adhesion molecule 1 (CEACAM1) has antiviral activity against human cytomegalovirus, influenza virus and metapneumovirus by supressing mTOR-mediated protein synthesis (33, 34). The membrane protein CEACAM1 is induced by innate sensors such as TLR-4 (35) and IFI16 (34) and delivers inhibitory signals via SHP1 (haematopoietic cells) or SHP2 (epithelial and endothelial cells) phosphatase activity through CEACAM1 immunoreceptor tyrosine-based inhibitory motifs (ITIMs) (36). CEACAM1 expression is rapidly induced following activation of NF-κB and IRF1, but whether IFN-α alone (as used here) can induce it expression is not clear. The *IRF1* promoter possesses a single GAS element, but no ISRE, and so its expression is induced by STAT1 homodimers (37). Type I IFN signalling predominantly leads to the formation of STAT1-STAT2 heterodimers that associate with IRF9 (to form the ISGF3 transcription factor) to drive expression of ISGs that possess ISRE elements in their promoters; however, STAT1 homodimers are formed after type I IFN treatment, but these are at lower concentrations. It is possible that ‘late’ inhibition of protein synthesis in ISG15-deficient cells (compared to the swifter antiviral activity of ISRE-containing genes such as IFIT1) may relate to the kinetics of CEACAM1 expression as the accumulation of STAT1 homodimers is required to drive the expression of *IRF1*, that itself needs to be translated before it induces *CEACAM1*. Of course, the accumulation of STAT1 homodimers may be higher in ISG15-deficient cells because of a dysregulated type I IFN response. Nevertheless, it is plausible that the overamplified type I IFN response in ISG15-deficient cells led to high levels of CEACAM1 (compared to control cells) resulting in inhibition of protein synthesis. Moreover, ISG15 may have yet-to-be characterised functions in regulating the cellular response to stressors that lead to inhibition of protein synthesis.

It has been reported that ISG15 has a role in regulating the cell cycle through its interactions with SKP2 and USP18, although experiments in that study were not performed in IFN-treated cells, nor were ISG15 knockout cells tested (15). While rates of protein synthesis differ during different stages of the cell cycle, translation is thought to be lowest during mitosis (38). Perturbation of the ISG15- SKP2-USP18 axis following ablation of USP18 led to a delayed progression from G1 to S phase which is not generally thought to be associated with translational repression (39). Of note, we have not observed any obvious differences in cell growth in non-treated A549-ISG15^-/-^ cells. Further work is required to dissect the mechanism responsible for ISG15’s effects on general protein translation during an antiviral response.

ISG15 has emerged as a central regulator of immunity. It is a pleotropic protein that is strongly expressed following activation of innate immune sensors and connects innate and adaptive immunity. In this study, we have shown that a lack of ISG15 leads to virus resistance by two kinetically distinct mechanisms; the rapid induction of antiviral ISGs and the unexpected effects on protein synthesis. Our newly developed cell lines and infection model will pave the way for further studies investigating the regulatory mechanisms of ISG15 during the antiviral response.

## Acknowledgments

We are grateful to ERASMUS+ for supporting DH. The authors also acknowledge technical support provided by Christopher Simmons-Riach and Miroslav Botev.

## Figure Legends

**Supplementary Figure 1.**
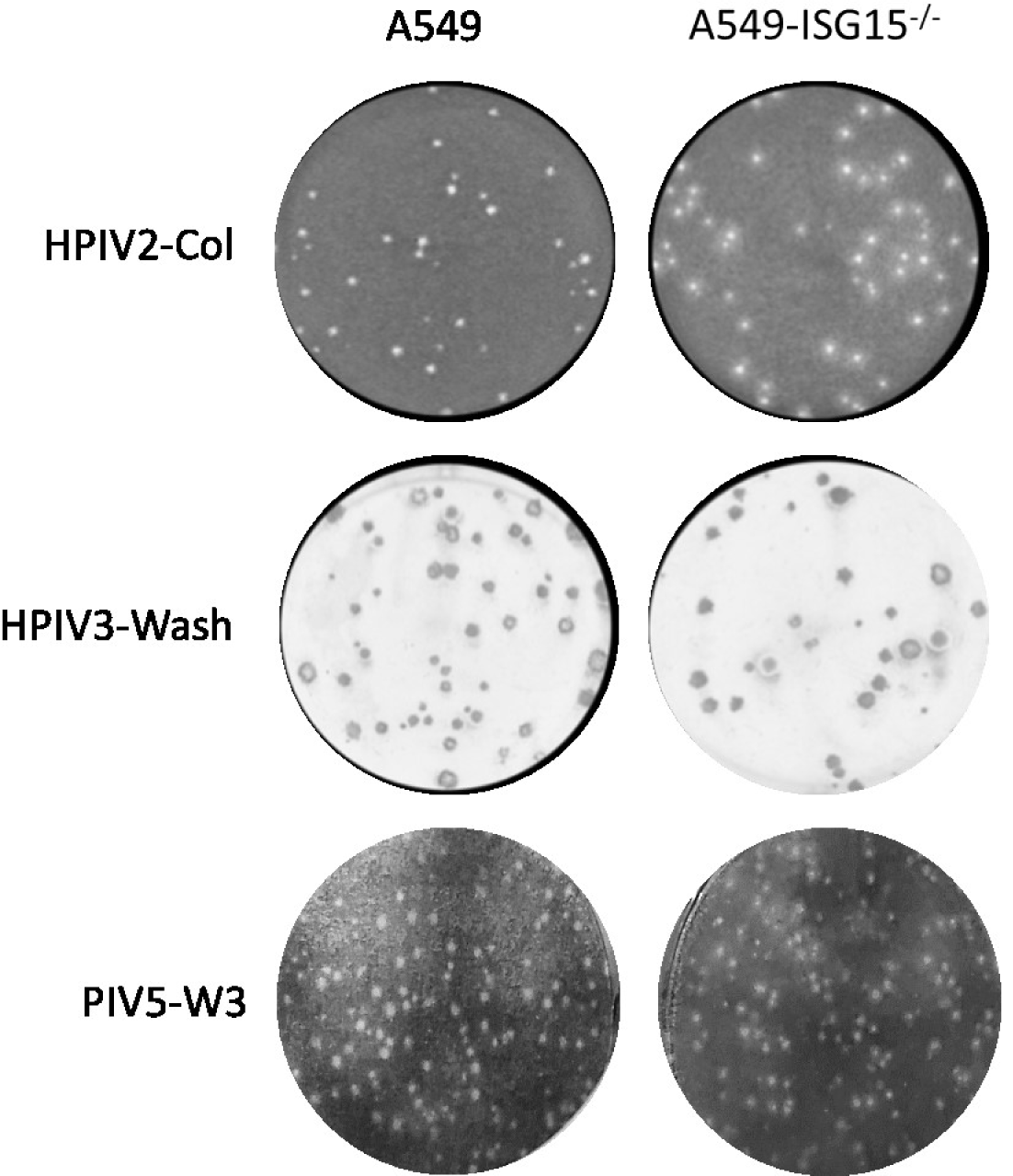
No viral resistance in naïve ISG15-deficient cells. Near confluent A549 or A549-ISG15^-/-^ (B8) cells in 6-well plates were infected with the indicated virus at dilution that allow the formation of discrete plaques. Following 6 days infection, cells were fixed and either stained with toluidine blue O (HPIV2 strain Collindale and PIV5 strain W3-infected cells) or immunostained (HPIV3 strain Washington using antibodies specific for HPIV3 nucleoprotein).

## Notes

### Competing Interest Statement

The authors have declared no competing interest.

## References

1. McFadden, M.J., N.S. Gokhale, and S.M. Horner. 2017. Protect this house: cytosolic sensing of viruses. Curr Opin Virol. 22: 36–43.

2. Kawai, T. and S. Akira. 2010. The role of pattern-recognition receptors in innate immunity: update on Toll-like receptors. Nat Immunol. 11(5): 373–384.

3. Randall, R.E. and S. Goodbourn. 2008. Interferons and viruses: an interplay between induction, signalling, antiviral responses and virus countermeasures. J Gen Virol. 89(Pt 1): 1–47.

4. Dalet, A., E. Gatti, and P. Pierre. 2015. Integration of PKR-dependent translation inhibition with innate immunity is required for a coordinated anti-viral response. FEBS Lett. 589(14): 1539–1545.

5. Mears, H.V. and T.R. Sweeney. 2018. Better together: the role of IFIT protein-protein interactions in the antiviral response. J Gen Virol. 99(11): 1463–1477.

6. Schoggins, J.W., S.J. Wilson, M. Panis, M.Y. Murphy, C.T. Jones, P. Bieniasz, and C.M. Rice. 2011. A diverse range of gene products are effectors of the type I interferon antiviral response. Nature. 472(7344): 481–485.

7. Yu, Z.X. and H.M. Song. 2019. Toward a better understanding of type I interferonopathies: a brief summary, update and beyond. World J Pediatr.

8. Perng, Y.C. and D.J. Lenschow. 2018. ISG15 in antiviral immunity and beyond. Nat Rev Microbiol. 16(7): 423–439.

9. Malakhov, M.P., O.A. Malakhova, K.I. Kim, K.J. Ritchie, and D.E. Zhang. 2002. UBP43 (USP18) specifically removes ISG15 from conjugated proteins. J Biol Chem. 277(12): 9976–9981.

10. Zhang, X., D. Bogunovic, B. Payelle-Brogard, V. Francois-Newton, S.D. Speer, C. Yuan, S. Volpi, Z. Li, O. Sanal, D. Mansouri, I. Tezcan, G.I. Rice, C. Chen, N. Mansouri, S.A. Mahdaviani, Y. Itan, B. Boisson, S. Okada, L. Zeng, X. Wang, H. Jiang, W. Liu, T. Han, D. Liu, T. Ma, B. Wang, M. Liu, J.Y. Liu, Q.K. Wang, D. Yalnizoglu, L. Radoshevich, G. Uze, P. Gros, F. Rozenberg, S.Y. Zhang, E. Jouanguy, J. Bustamante, A. Garcia-Sastre, L. Abel, P. Lebon, L.D. Notarangelo, Y.J. Crow, S. Boisson-Dupuis, J.L. Casanova, and S. Pellegrini. 2015. Human intracellular ISG15 prevents interferon-alpha/beta over-amplification and auto-inflammation. Nature. 517(7532): 89–93.

11. Francois-Newton, V., G. Magno de Freitas Almeida, B. Payelle-Brogard, D. Monneron, L. Pichard-Garcia, J. Piehler, S. Pellegrini, and G. Uze. 2011. USP18-based negative feedback control is induced by type I and type III interferons and specifically inactivates interferon alpha response. PLoS One. 6(7): e22200.

12. Malakhova, O.A., K.I. Kim, J.K. Luo, W. Zou, K.G. Kumar, S.Y. Fuchs, K. Shuai, and D.E. Zhang. 2006. UBP43 is a novel regulator of interferon signaling independent of its ISG15 isopeptidase activity. EMBO J. 25(11): 2358–2367.

13. Sarasin-Filipowicz, M., X. Wang, M. Yan, F.H. Duong, V. Poli, D.J. Hilton, D.E. Zhang, and M.H. Heim. 2009. Alpha interferon induces long-lasting refractoriness of JAK-STAT signaling in the mouse liver through induction of USP18/UBP43. Mol Cell Biol. 29(17): 4841–4851.

14. Speer, S.D., Z. Li, S. Buta, B. Payelle-Brogard, L. Qian, F. Vigant, E. Rubino, T.J. Gardner, T. Wedeking, M. Hermann, J. Duehr, O. Sanal, I. Tezcan, N. Mansouri, P. Tabarsi, D. Mansouri, V. Francois-Newton, C.F. Daussy, M.R. Rodriguez, D.J. Lenschow, A.N. Freiberg, D. Tortorella, J. Piehler, B. Lee, A. Garcia-Sastre, S. Pellegrini, and D. Bogunovic. 2016. ISG15 deficiency and increased viral resistance in humans but not mice. Nat Commun. 7: 11496.

15. Vuillier, F., Z. Li, P.H. Commere, L.T. Dynesen, and S. Pellegrini. 2019. USP18 and ISG15 coordinately impact on SKP2 and cell cycle progression. Sci Rep. 9(1): 4066.

16. Hermann, M. and D. Bogunovic. 2017. ISG15: In Sickness and in Health. Trends Immunol. 38(2): 79–93.

17. Andrejeva, J., H. Norsted, M. Habjan, V. Thiel, S. Goodbourn, and R.E. Randall. 2013. ISG56/IFIT1 is primarily responsible for interferon-induced changes to patterns of parainfluenza virus type 5 transcription and protein synthesis. J Gen Virol. 94(Pt 1): 59–68.

18. Domingues, P., C.G. Bamford, C. Boutell, and J. McLauchlan. 2015. Inhibition of hepatitis C virus RNA replication by ISG15 does not require its conjugation to protein substrates by the HERC5 E3 ligase. J Gen Virol. 96(11): 3236–3242.

19. Sanjana, N.E., O. Shalem, and F. Zhang. 2014. Improved vectors and genome-wide libraries for CRISPR screening. Nat Methods. 11(8): 783–784.

20. Durbin, A.P., J.M. McAuliffe, P.L. Collins, and B.R. Murphy. 1999. Mutations in the C, D, and V open reading frames of human parainfluenza virus type 3 attenuate replication in rodents and primates. Virology. 261(2): 319–330.

21. Hilton, L., K. Moganeradj, G. Zhang, Y.H. Chen, R.E. Randall, J.W. McCauley, and S. Goodbourn. 2006. The NPro product of bovine viral diarrhea virus inhibits DNA binding by interferon regulatory factor 3 and targets it for proteasomal degradation. J Virol. 80(23): 11723–11732.

22. Choppin, P.W. 1964. Multiplication of a Myxovirus (Sv5) with Minimal Cytopathic Effects and without Interference. Virology. 23: 224–233.

23. Didcock, L., D.F. Young, S. Goodbourn, and R.E. Randall. 1999. The V protein of simian virus 5 inhibits interferon signalling by targeting STAT1 for proteasome-mediated degradation. J Virol. 73(12): 9928–9933.

24. Randall, R.E., D.F. Young, K.K. Goswami, and W.C. Russell. 1987. Isolation and characterization of monoclonal antibodies to simian virus 5 and their use in revealing antigenic differences between human, canine and simian isolates. J Gen Virol. 68 (Pt 11): 2769–2780.

25. Rydbeck, R., C. Orvell, A. Love, and E. Norrby. 1986. Characterization of four parainfluenza virus type 3 proteins by use of monoclonal antibodies. J Gen Virol. 67 (Pt 8): 1531–1542.

26. Wignall-Fleming, E.B., D.J. Hughes, S. Vattipally, S. Modha, S. Goodbourn, A.J. Davison, and R.E. Randall. 2019. Analysis of Paramyxovirus Transcription and Replication by High-Throughput Sequencing. J Virol. 93(17).

27. Young, D.F., J. Andrejeva, X. Li, F. Inesta-Vaquera, C. Dong, V.H. Cowling, S. Goodbourn, and R.E. Randall. 2016. Human IFIT1 Inhibits mRNA Translation of Rubulaviruses but Not Other Members of the Paramyxoviridae Family. J Virol. 90(20): 9446–9456.

28. Killip, M.J., D.F. Young, C.S. Ross, S. Chen, S. Goodbourn, and R.E. Randall. 2011. Failure to activate the IFN-beta promoter by a paramyxovirus lacking an interferon antagonist. Virology. 415(1): 39–46.

29. Killip, M.J., D.F. Young, D. Gatherer, C.S. Ross, J.A. Short, A.J. Davison, S. Goodbourn, and R.E. Randall. 2013. Deep sequencing analysis of defective genomes of parainfluenza virus 5 and their role in interferon induction. J Virol. 87(9): 4798–4807.

30. Chen, C., R.W. Compans, and P.W. Choppin. 1971. Parainfluenza virus surface projections: glycoproteins with haemagglutinin and neuraminidase activities. J Gen Virol. 11(1): 53–58.

31. Young, D.F., E.B. Wignall-Fleming, D.C. Busse, M.J. Pickin, J. Hankinson, E.M. Randall, A. Tavendale, A.J. Davison, D. Lamont, J.S. Tregoning, S. Goodbourn, and R.E. Randall. 2019. The switch between acute and persistent paramyxovirus infection caused by single amino acid substitutions in the RNA polymerase P subunit. PLoS Pathog. 15(2): e1007561.

32. Bogunovic, D., M. Byun, L.A. Durfee, A. Abhyankar, O. Sanal, D. Mansouri, S. Salem, I. Radovanovic, A.V. Grant, P. Adimi, N. Mansouri, S. Okada, V.L. Bryant, X.F. Kong, A. Kreins, M.M. Velez, B. Boisson, S. Khalilzadeh, U. Ozcelik, I.A. Darazam, J.W. Schoggins, C.M. Rice, S. Al-Muhsen, M. Behr, G. Vogt, A. Puel, J. Bustamante, P. Gros, J.M. Huibregtse, L. Abel, S. Boisson-Dupuis, and J.L. Casanova. 2012. Mycobacterial disease and impaired IFN-gamma immunity in humans with inherited ISG15 deficiency. Science. 337(6102): 1684–1688.

33. Diab, M., A. Vitenshtein, Y. Drori, R. Yamin, O. Danziger, R. Zamostiano, M. Mandelboim, E. Bacharach, and O. Mandelboim. 2016. Suppression of human metapneumovirus (HMPV) infection by the innate sensing gene CEACAM1. Oncotarget. 7(41): 66468–66479.

34. Vitenshtein, A., Y. Weisblum, S. Hauka, A. Halenius, E. Oiknine-Djian, P. Tsukerman, Y. Bauman, Y. Bar-On, N. Stern-Ginossar, J. Enk, R. Ortenberg, J. Tai, G. Markel, R.S. Blumberg, H. Hengel, S. Jonjic, D.G. Wolf, H. Adler, R. Kammerer, and O. Mandelboim. 2016. CEACAM1-Mediated Inhibition of Virus Production. Cell Rep. 15(11): 2331–2339.

35. Muenzner, P., M. Naumann, T.F. Meyer, and S.D. Gray-Owen. 2001. Pathogenic Neisseria trigger expression of their carcinoembryonic antigen-related cellular adhesion molecule 1 (CEACAM1; previously CD66a) receptor on primary endothelial cells by activating the immediate early response transcription factor, nuclear factor-kappaB. J Biol Chem. 276(26): 24331–24340.

36. Gray-Owen, S.D. and R.S. Blumberg. 2006. CEACAM1: contact-dependent control of immunity. Nat Rev Immunol. 6(6): 433–446.

37. Michalska, A., K. Blaszczyk, J. Wesoly, and H.A.R. Bluyssen. 2018. A Positive Feedback Amplifier Circuit That Regulates Interferon (IFN)-Stimulated Gene Expression and Controls Type I and Type II IFN Responses. Front Immunol. 9: 1135.

38. Polymenis, M. and R. Aramayo. 2015. Translate to divide: small es, Cyrillicontrol of the cell cycle by protein synthesis. Microb Cell. 2(4): 94–104.

39. Stumpf, C.R., M.V. Moreno, A.B. Olshen, B.S. Taylor, and D. Ruggero. 2013. The translational landscape of the mammalian cell cycle. Mol Cell. 52(4): 574–582.

